# Design and development of online pressure sensing for microbial batch cultivation

**DOI:** 10.64898/2026.06.08.729494

**Authors:** Martin Malthe Borch, Pauline Kehr, Rafael Altamirano Torres, Philip J. Gorter de Vries, Niels Jakob Larsen, Nicolas Padfield, Alex Toftgaard Nielsen

## Abstract

Gas production and consumption is a direct consequence of microbial activity in environmental and industrial settings. In closed batch cultivations, headspace pressure changes therefore give valuable insights into the microbial metabolism. For laboratory scale anaerobic batch cultivations, manual manometer measurements are routinely applied, as a simple and robust method, but it is labour intensive, causes disturbances in the headspace gas and temperature, leading to suboptimal growth, inhibition and noisy data. We built and tested an automated online pressure sensor for closed batch cultivations. It is designed for microbial cultivation and integrates with sterile and anaerobic cultivation workflows. The system uses an absolute pressure sensor (0-30 bar) mounted on a custom designed PCB, with a gas-tight needle mount. An ESP32 microcontroller logs pressure and temperature locally and generates a Wi-Fi access point for real-time visualization and direct CSV download through a local homepage. We detail hardware and software design decisions, assembly, and validation including long-term stability. Case studies demonstrate the applicability for: a multiphasic biogas kinetics during anaerobic digestion, capturing gas uptake dynamics and metabolic shifts during syngas fermentations and co-feeding experiments, and long-term robustness in a multi-year monitoring of a compressed-air system. More than 130 individual sensors have been deployed over 3 years in laboratories, at various academic and industrial settings. The platform provides reproducible, high-resolution pressure measurements that enable calculation of gas formation/consumption rates and improve experimental throughput without disturbing cultures. Design files, firmware, and example analysis scripts are openly available to support adoption and further development.

## 1 Introduction

Gas production or consumption is central in microbial metabolism and fundamental to the environmental cycles of carbon (CO, CO_2_, CH_4_), oxygen (O_2_), hydrogen (H_2_), sulfur (H_2_S), and nitrogen through denitrification processes (N_2_, N_2_O, NO) (Banfield and Nealson 1997; Beulig et al. 2019; Wang et al. 2011). These processes are also widespread in industrial biotechnological processes, such as wastewater treatment, biogas, or bioethanol production. Measuring soil microbial respiration (CO_2_) (Stotzky 1965) is fundamental in environmental and agricultural research, and several techniques are available (Freijer and Bouten 1991; Bond-Lamberty et al. 2024). Lastly, and familiar to most, CO2 is generated during food fermentation, including brewing, yoghurt, cheese, sauerkraut, and baking. With an ongoing focus on mitigating climate change and CO_2_ emissions (Mukherji 2023), bacteria have received increased attention, as they can consume gases such as CO_2_, H_2_, and CO, with applications in the production of sustainable chemicals and biomanufactured materials, have (Redl et al. 2017).

The microbial activity can be divided into catabolism, the energy-conserving reactions, and anabolism, the reactions leading to the formation of biomass (Villadsen et al. 2011). The formation or consumption of gas occurs mainly during catabolic reactions. Detection of threshold levels of gas pressure also marks the point at which microbial activity stops, and the energy driving (thermodynamic) reactions is balanced. This can be used to characterise the microbes and calculate the energy required for growth, or generated in relation to the products produced (ATP mol/product) (Laura and Jo 2023; Philips 2020). Historically, light has been used to measure the activity of bacteria, increasing light absorption, which equals the growth, and is referred to as optical density (OD) (Koch 1961; Pasteur 1857). But this method only represents one half of the metabolism, the anabolic reactions, and it is impossible to get reliable OD measurements for biofilms, solid state fermentation, or when the medium is black or filled with particles. Also, OD doesn’t measure microbial activity in the absence of growth. In closed-batch cultivations, undisturbed, high-quality online pressure measurements enable the characterisation and assessment of microbial catabolic activity, which is particularly relevant when OD cannot be used, or no growth is occurring.

The standard anaerobic laboratory cultivation method uses flasks or vials with butyl-rubber stoppers, enabling needle-based manipulation and sampling (Mauerhofer et al. 2019; Hanišáková et al. 2022). For these cultivations, pressure is used qualitatively to evaluate gas change reactions by measuring the pressure in the closed headspace. Although automated measurements and plate methods have been described (Nakamura et al. 2011), and plate readers have been integrated into large anaerobic chambers (Clavel et al. 2025), manual sampling remains widely used. This is due to cost, the difficulty of combining electronics with explosive gases such as H_2_ and CO, gas leakage at high pressure, particularly H_2_, and the challenges of maintaining sterile, anaerobic conditions.

Although manual sampling is commonly used, it presents several challenges that also motivated and initiated this work, as they were encountered during ongoing research on gas fermentation. A series of sequential batch cultivations using syngas was performed to enrich gas-consuming microbes (Borch et al. 2026). The initial slurry was black, making optical measurements of biomass challenging. Pressure sampling with a manual manometer was employed, but several issues arose. Moving the vials out of the incubator caused temperature fluctuations, leading to pressure variations that were difficult to control and resulting in drops in incubator temperature during sampling. The initial growth was limited and extremely slow, and the pressure decrease correlated with both the sampling rate and the growth rate. Each sample could cause a pressure drop of 10-50 mbar, depending on the total pressure, headspace volume, and dead volume of the sampling device. Limited sampling was necessary but led to poor monitoring, causing the cultures to enter the stationary phase and sporulate, thereby missing the optimal time point for subcultivation. It was also time-consuming, as there were five conditions at three temperatures, each in duplicate, a total of 45 vials. Although ultimately successful, the manual sampling process extended the experiment’s overall duration and yielded suboptimal data. These specific challenges, and the reporting of similar challenges for biogas potentials (Himanshu et al. 2017; Pérez-Vidal et al. 2025; Fdz-Polanco et al. 2005) highlighted the limitations of manual sampling and prompted an investigation into other possibilities.

Current pressure sensor solutions can broadly be divided into three categories: (1) commercially available laboratory solutions, (2) high-quality sensors made for industrial fermentations and chemical installations, and (3) various commercially available sensor breakout boards for prototyping. Commercially available laboratory sensors are primarily designed to measure biogas potential, CO2 formation during food fermentation, or oxygen demand in aerobic fermentations. They generally come as integrated platforms for fermentation and work exceedingly well for those applications, (Fdz-Polanco et al. 2005; Pérez-Vidal et al. 2025; Himanshu et al. 2017), but are challenging to repurpose more broadly. The available solutions were costly, did not fit the sterile, anaerobic vial cultivation setup with rubber stoppers, and had a pressure range that was too low for our needs. Industrial sensors are designed for specific software integration and hardware compatibility and are often also approved for use in applications involving explosive gases (ATEX). They tend to be large, costly, and overly specific for laboratory use. Lastly, the so-called breakout boards are typically small surface-mounted device (SMD) components mounted on a custom Printed Circuit Board (PCB). Though plenty available, they were not designed to allow for pressure-tight needle connections. For lists and comparison, see the supplementary materials.

In recent years, open-source software and hardware movements like the Arduino project, the open-source hardware association (OSHWA), and the hackerspace movements (Davies 2017), have broadened to include biology and biological engineering, in movements such as Biohackers and DIYBio (Delfanti 2013; Delgado 2013). Some recent open source hardware laboratory equipment projects include: clamp-on and real-time photometers for microbial cultivations (Deutzmann et al. 2022), qByte for nucleic acids and protein analysis (Quero et al. 2025), an electroporator (Cauchy et al. 2023), a DSLR-based plate colony analyser (Pi et al. 2022), and NanoMi an open source electron microscope platform (Malac et al. 2022). Such prototyping projects can be supported by FabLabs, either independent or university-based environments that provide tools and education for digital fabrication and rapid prototyping to students (Svabo and Borch 2020), researchers, and the general public (Padfield et al. 2014; Haldrup et al. 2018).

The aim of this work was to develop an automated online pressure-sensing system that enables high-sensitivity, high-precision measurements without disturbing the experimental setup. To gain new insights into microbial catalytic activity through gas uptake or production rates, information that cannot be obtained from manual pressure sampling in standard small-scale vial experiments or from OD measurements. We document the design process, key challenges, and innovations involved in developing this pressure sensor system. The parts are: 1) design criteria and identification of a suitable sensor, 2) description of hardware and software solutions, 3) reporting on case applications within biogas production and gas fermentation, and 4) discussing the method, results, and perspectives.

## 2 Materials and Equipment

### 2.1 List of materials

The built system comprises the following components and materials. Details of each component are listed below.

- Printed circuit board (PCB) designed by us but produced by a commercial manufacturer. It has a pressure sensor, the MS5837-30BA from TE connectivity mounted on it.
- Code.
- Arduino IDE (Integrated Development Environment) software.
- Microcontroller Wemos ESP32 on the D1 mini prototyping board.
- Needle holder, laser-cut 4 mm acrylic, stainless steel, or PCB material, and bolts.
- Gasket. Butyl rubber, laser cut.
- A microSD-card converter soldered to the ESP32.
- A microSD-card.

### 2.2 Material details

Printed circuit board (PCB) designed by us, but produced by a commercial manufacturer (JLCpcb). The design files are available at github.com/nxdf/laerke. This includes the design files for the PCB, needle holder, rubber gasket, and source code.

The sensor PCB included the following components and features: a pressure sensor (MS5837-30BA) from TE Connectivity. This is a 0-30 bar absolute (not relative) sensor with a resolution of 0.2 mBar and a claimed accuracy of +/- 1.5 mBar (if calibrated), +/- 50 mBar otherwise. We observed differences of the order of 50 mBar between sensors, that was corrected during data analysis. Capacitor 100 nF for power supply decoupling. Two resistors, 10kΩ for data and clock pull-up, are designed with solder pads for easy enabling and disabling. The data line pull-up resistor is default enabled, and the clock line pull-up is default disabled. As the clock is shared between all sensors, there should be only one clock pull-up resistor for the whole clock bus, not one per sensor. If one mounts, e.g., 4 sensors to one microcontroller, one must remember to bridge the clock pull-up resistor solder pads on one of the sensors (but only one). The data line pull-up has solder pads; these are already bridged in the circuit board design. It is possible to cut the trace with a knife, but this is not normally necessary; it is a feature provided only for experiments or unusual wiring setups. Wire connections: There is space for a standard 2.54 mm pin header, a Qwiic connector (for wires with plugs), or soldering of wires directly to the pads. Four conductors are required: power (3.3V), ground, I2C data and I2C clock. PCB labelling of pads, connection pins, a circle to help position the rubber gasket, the sensor part number and I2C address (0×76).

Code. The code is written in C for Arduino, open source, and available at github.com/nxdf/laerke. Arduino IDE (Integrated Development Environment) software was used for coding, compiling, and interfacing.

Microcontroller Wemos ESP32 on the D1 mini prototyping board. Most other ESP32 boards would work. The board handles computing, sensor input, data sharing, SD card, and Wi-Fi hotspot. Depending on the board used, it will have about 26-34 I/O pins, of which 4 are required for interfacing with the SD card, one is required for the I2C clock line, and one is used to power the sensors so they can be reset, increasing reliability. This leaves about 20-28 i/o pins for software I2C, so it should be possible to have up to about 20 sensors per board. We have tested 8. We selected the ESP32 because it has Wi-Fi and is affordable; any microcontroller could be used if Wi-Fi is not required.

Needle holder, laser cut 4 mm acrylic/stainless steel/PCB (printed circuit board) material. We tested various materials – laser cut acrylic, laser cut stainless steel, and having the PCB (printed circuit board) manufacturer produce the holder out of PCB material (which is glass fibre reinforced plastic). All seemed to work fine. The two halves of the needle holder compress the needle against the rubber seal using two M4 bolts.

Gasket. Butyl rubber, laser cut. These were cut in the laser between two layers of cardboard to keep them in place. Note that the gasket seals to the PCB, not only to the sensor. We found it difficult to achieve a good seal if sealing only to the sensor itself.

A microSD-card converter soldered to the ESP32. An SD card module would work equally well. A microSD-card.

## 3 Methods

### 3.1 Design criteria

We initially defined a series of design criteria to guide the design phase and ensure the desired outcome. The following points were defined:

- Absolute pressure sensing from below atmospheric to + 20 bar.
- Integration with existing sterile anaerobic workflow with vials and needles. Gastight enough for use with hydrogen.
- Possible to apply with explosive gases (ATEX).
- Inexpensive, making it economically feasible to measure dozens of vial experiments.
- Data logging, backup, and visualization.
- The simplest, easiest, and fastest interface and user flow for mounting, dismounting, starting the experiment, and initiating logging.
- Robustness and simplicity take priority over features and modularity.

### 3.2 Final design

The design is based on the sensor breakout boards solution as mentioned above. The final system features an ESP32 microcontroller, three or more pressure sensors, and an SD card for local data storage. Initial system settings are configured through a config file on the SD card, which simplifies hardware requirements and the user interface while ensuring data consistency across experiments. When powered on, the sensor begins logging data for a new experiment. The microcontroller creates a Wi-Fi access point using the SSID and password stored in the SD card’s configuration file. It hosts a webpage that lets users view live sensor data, download data as a CSV file, or download a log file.

### 3.3 Hardware design process and decisions

Several factors affected our design and construction process: A breakout board-style design. Our solution fits into the category of breakout boards described above but features a form factor tailored for a specific use case. Modern sensors used in commercial electronics are typically small SMD (surface-mounted device) components, which are difficult to solder wires to. Therefore, a common approach in DIY, experimental, and rapid prototyping fields is to design and produce a small PCB (Printed Circuit Board) whose sole purpose is to mount the sensor component and provide larger electrical connectors. These breakout boards often include a few supporting components for the sensor, such as pull-up or pull-down resistors, smoothing capacitors, and power supply components. The commercially available boards examined did not leave enough space around the sensor itself for an airtight O-ring or gasket mount, nor did they have attachment points or holes for through-bolt mounting to properly compress a seal. By designing and producing our own board, we halved the cost compared to commercially available options, which is important when numerous sensors are needed. The design approach also allowed testing various configurations, soldering methods, needle connectors, and solutions to improve gas-tightness before finalizing the design.

Sensor Selection. The sensor was chosen after reviewing options from major global electronic component suppliers to ensure it was readily available, affordable, and had a reliable supply. The selected model is the MS5837-30BA from TE Connectivity, a high-precision sensor used in diving and underwater robot applications. It communicates via digital protocols (I2C), and we utilized an existing Arduino protocol and library that simplifies software development for communication and data logging. The sensor also offers temperature measurements, which are useful for calculating or correcting pressure readings affected by temperature changes. Key features include: factory calibration, an operating absolute pressure range from 0 to 30 bar, a maximum pressure of 50 bar, digital readings every 50-100 ms, a pressure resolution of 0.2 mbar, and a low error rate at extreme temperatures and pressures. It operates within a temperature range of −20 to +85°C. For a full description, see the manufacturer’s datasheet.

Gastight needle connection. This was optimized because H_2_ can leak through plastics and the needle head itself. After testing various needle-mounting solutions over several months, the final design directly attaches the needle to the PCB. Although this required redesign, it was essential to ensure hydrogen tightness by addressing issues with sliding or ill-fitting needle holders, leaking fittings, and gaskets. This was critical since experiments could last up to several months. The design also allows for sterile needle replacement. Many other pressure sensor solutions rely on tubing, which is another source of vulnerability to hydrogen leaks or oxygen ingress. To prevent this, no tubing or connectors were used. Some breakout boards with hose-attachment ports, common in recent research, were avoided because hoses increase the number of seals and the potential for leaks. Silicone tubing, in particular, is highly permeable to oxygen.

### 3.4 Manufacturing and assembly

The printed circuit board design was created in Fusion 360 (Figure 2), and components mounted during fabrication. For the PCB design and wiring, see the diagram Figure 2. The design files can be found at github.com/nxdf/laerke. The wires and the SD card holder were hand-soldered. Assembly and mounting of the sensor board to a sterile needle are done using M4 bolts and nuts.

**Figure 1.**
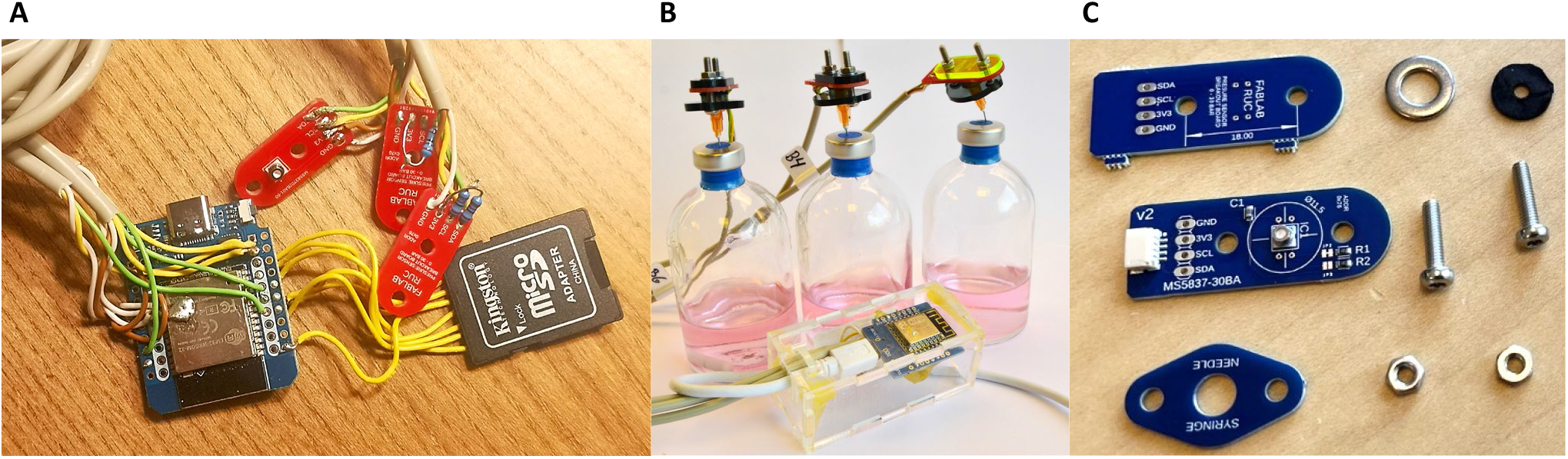
Pressure sensors. A) Pressure sensor system with 3 sensor PCBs and SDcard. B) Early prototype without SDcard and sensors mounted in vials. C) Parts to assemble a single senor, PCB with 4 pin I2C connector.

**Figure 2.**
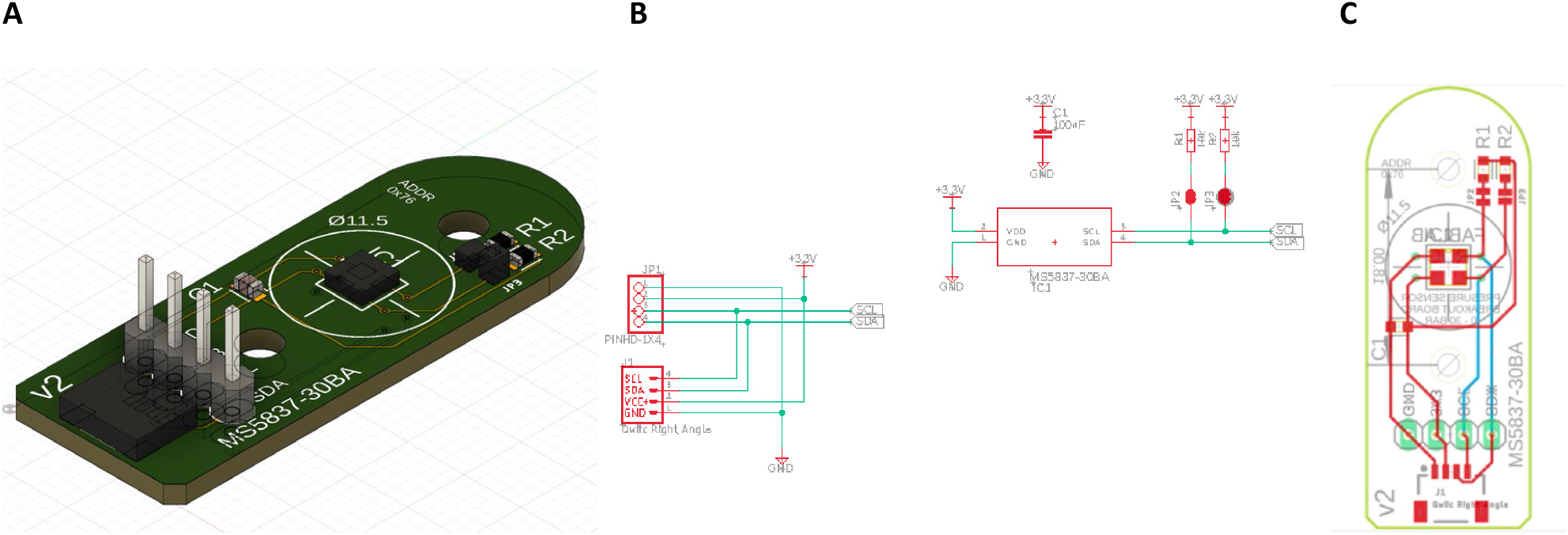
A) A 3D model of the assembled sensor board, generated in Fusion 360. B) Circuit diagram including capacitor and resistor values (100nF and 10kΩ). C) A view of the sensor board showing copper traces on both sides and silk screen printing simultaneously. High-resolution design files are available on GitHub github.com/nxdf/laerke.

### 3.5 Code and software design process and decisions

The microcontroller firmware was implemented in C and compiled using the Arduino IDE. The software contains different code blocks needed to run the system. There is the main loop running on the microcontroller and the libraries needed for communication with the pressure sensor. A key code element is a break switch loop “Software I2C multiplexing”, used to individually reach each sensor. This allows dynamically switching I2C data lines while sharing a common clock line. Before reading each new sensor, it was necessary to call Wire.end() and Wire.begin() with each pin number. The authors of the I2C library may not have envisioned this use case of changing the pin number before each reading. Description of each code element is in the supplementary materials, the full code and details can be found on github github.com/nxdf/laerke.

### 3.6 Initial validation and hydrogen permeability

We tested the long-term stability of the system for 1.5 months with 1.5 bars overpressure of atmospheric air. No pressure loss was detected in this experiment, and pressure fluctuations were attributed to room temperature fluctuations. (Supplementary materials). To measure hydrogen permeability, we compared the headspace of N_2_:CO_2_ (80:20) and H_2_:CO_2_ (80:20) mixtures at an overpressure of 1.7 bars at 57 °C. The N_2_:CO_2_ pressure dropped 0.2% in 3 days and the H_2_:CO_2_ 1.7% in 3 days. When accounting for the control and the hydrogen fraction, we find that hydrogen is leaking with 0.4% per day (Supplementary materials). For experiments with extended duration and a high hydrogen partial pressure p(H_2_), this should be taken into consideration in the experimental design. If needed, adding control and blank conditions can account for this loss during the analysis. This was considered an acceptable rate for the further case validation experiments.

### 3.7 Reproducibility and robustness

As of now, over 130 individual sensors have been utilised in laboratory and workshop experiments. They have been employed by three companies and ten academic research laboratories across various countries and continents, for a wide range of uses. The sensors have been used to cultivate thermophilic bacteria in incubators set to 60 °C. The biogas experiment below ran for 50 days (Results Section 4.1), and the sensor has been used continuously and reliably for three years for monitoring pressure of e.g. compressed air systems at Fablab RUC (Results Section 4.4). The following case studies further illustrate the quality and reproducibility of the experimental data.

## 4 Results

### 4.1 Biogas rates and production in two phases

One of the most successful biotechnological processes is biogas production with +21.000 operational biogas and biomethane plants operational in Europe alone according to (The European Biogas Association 2025). Under industrially relevant conditions, the medium is black and contains suspended solids (Li et al. 2025), making it impossible to use spectrophotometric methods for growth estimations. Pressure measurements, combined with gas-composition sampling, are routine in vial experiments screening anaerobic digestion conditions (Velasquez-Pinas et al. 2025). We validated the online pressure sensors for monitoring and analysing biogas formation in standard vial experiments. We estimated the biogas potential of cattle manure, compared with a cellulose control and a blank containing only the inoculum. Cultivation was performed in standard 120 ml anaerobic crimp vials with butyl rubber stoppers, and the vials were incubated at 37°C for 50 days. The data was analysed and plotted in R (R file available). In the plot of the raw pressure sampling points for the three conditions, several small pressure drops are visible due to temperature fluctuations, as the incubator was frequently opened (Figure 3). The average temperature of all the sensors during the experiment was 37.08 °C; thus, the observed temperature drops were not an experimental issue. The duplicate conditions align perfectly, creating continuous lines without any data fitting or smoothing required.

**Figure 3.**
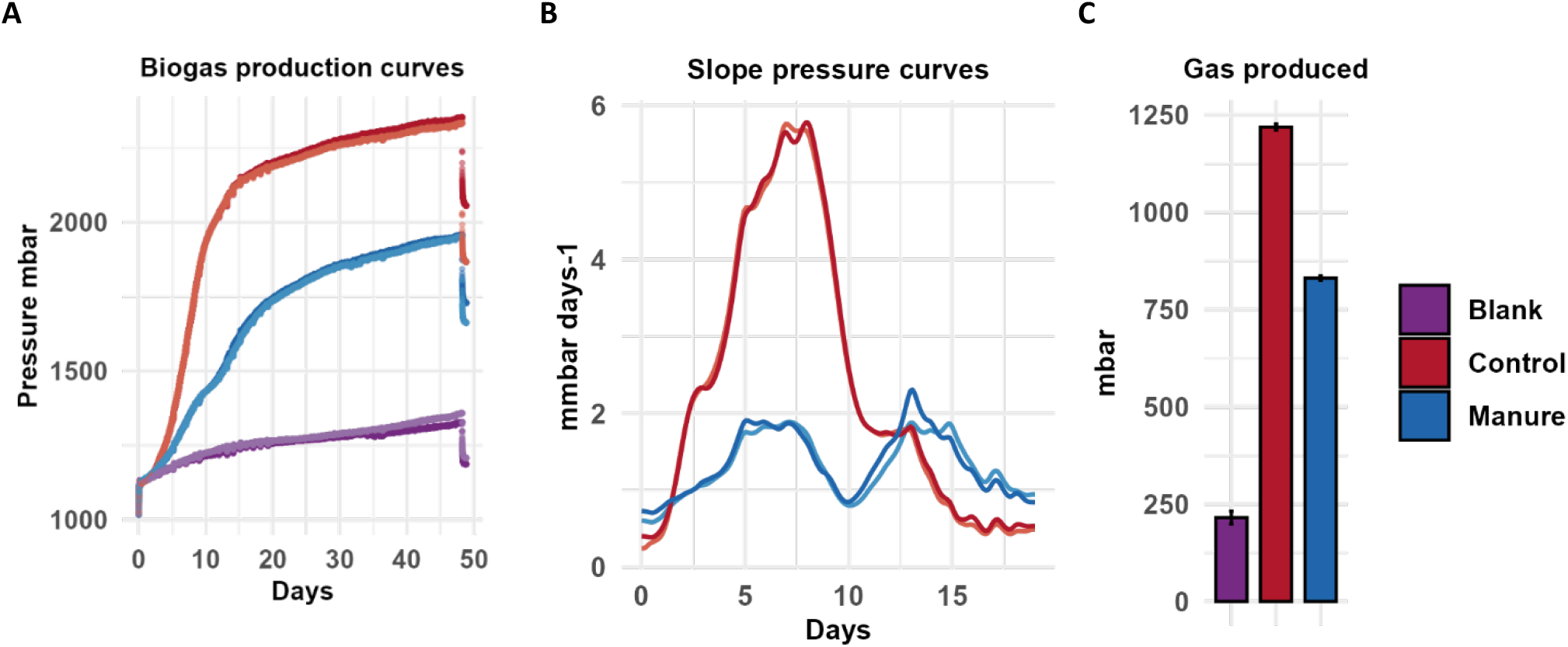
Biogas formation in anaerobic digestion. A) Pressure curves with gas formation in mbar. B) Gas production rate (slope) for the control (cellulose) and manure conditions. C) Bar plot of produced gas as increase in mbar. Errorbars are standard error of duplicates.

Calculating the gas formation. We inserted the pressure sensors before placing the vials in the incubator. The pressure increases from room temperature at 23 °C to 37 °C was, for all the vials, an average of 107.6 mbar. This aligns well with a pressure increase due to the water vapor pressure (62.8 mbar) and a dry gas pressure increase from 23 to 37 °C (47.6 mbar), a total of 110.4 mbar. This initial pressure increase was subtracted from the pressure increase to calculate gas production. The gas production was 215, 831, and 1219 mbar for the blank, manure, and control, respectively. With a vial of 120 ml and a liquid volume of 50 ml, and subtracting the blank, that gives a gas production of 1.67 and 2.72 mmol gas produced for the manure and control conditions. The gas formation rate was obtained as the slope of a spline fit. Two phases were detected in cattle manure digestion, peaking at about the same rate after 5 and again after approx. 12.5 days, each with a duration of about 2.5 days. We expect the first phase to be degradation of suspended carbohydrates and fatty acids, and the second phase the results of the microbial hydrolysis of the lignocellulosic materials in the manure (DelaVega-Quintero et al. 2025). The two identified peaks in the gas rate show the potential of the sensors to reveal the fermentation kinetics of complex substrates.

The stability and the efficiency of a biogas process can be increased by applying a two-stage fermentation process, taking into account these different phases (Shoukat et al. 2025), and therefore efforts are made to model the kinetics of the biogas production (Ali et al. 2021). Recently biochar addition to anaerobic biogas batch experiments has been shown to “dramatically” shorten the lag phase and increase the gas production rate, due to efficient, direct electron transfer between syntrophic fatty acid oxidizers and methanogens (Li et al. 2018; Velasquez-Pinas et al. 2025). Uninterrupted online pressure sampling could help further clarify the kinetic effects of complex substrates, as well as the impact of additions and co-feeding strategies. This can again support model development, process design, monitoring, and optimization of biogas systems.

**Table 1.** Anaerobic digestion results. The measured pressure increase from the anaerobic digestion cultivations.

| Condition | Gas production |  | Corrected |  |
| --- | --- | --- | --- | --- |
|  | mbar | mmol | mbar | mmol |
| Blank | 215 | 0.59 |  |  |
| Manure | 831 | 2.26 | 616 | 1.67 |
| Control | 1219 | 3.31 | 1004 | 2.72 |

### 4.2 Co-feeding kinetics and parallel metabolism in mixed community syngas fermentations

In gas-fermentation processes, gases such as CO_2_, H_2_, and CO provide both energy and a carbon source for microbial growth; thus, CO_2_ is consumed, unlike in sugar-based fermentations, where CO_2_ is produced. One of the best-known and most efficient microbial carbon-fixating pathways is the Wood-Ljungdahl pathway (WLP). H_2_ provides the reducing energy and donates electrons to CO2, converting it into acetyl-CoA and then acetate. The processes do not generate ATP through substrate-level phosphorylation, and the microbes therefore depend on energy generation from proton influx through an ATPase complex (Schuchmann and Müller 2014). The use of gaseous feedstocks, though, causes several challenges. The low solubility of gases, particularly hydrogen and carbon monoxide, limits gas mass transfer into the reaction medium (Gorter de Vries et al., 2024), and the limited energy available in the process results in an ATP-generation bottleneck for biomass formation. Co-feeding strategies were shown to enhance gas fermentation performance. Adding fructose to a *Moorella thermoacetica* H_2_:CO_2_ fermentation increased carbon fixation rates, with an optimal supplementation of 9% electrons from sugars. Beyond this level, metabolism shifted from gas fermentation to sugar fermentation (Park et al. 2019). Naturally monitoring pressure in closed batch gas fermentation is essential to determine if and how quickly gases are consumed by microbes and converted into liquid products.

Co-feeding strategies were investigated to determine how various approaches could improve the efficiency of thermophilic syngas fermentations, with the aim of investigating sustainable chemicals production. Fructose and a lignocellulosic hydrolysate were co-fed to a microbial community and a pure culture of *M. thermoactica* at various amounts and time points. The microbial community was a new enrichment made for this work, with a previously described methodology (Borch et al. 2026). The gas pressure response was analysed using the online pressure sensors. Specifically, fructose was co-fed to syngas fermentation at various time points, and the results were compared with cultivations fed only fructose or syngas.

With fructose alone, gases were produced rapidly and then stopped sharply. The pressure then stayed constant until a gradual and increasing uptake was seen (Figure 4A). This aligns with our earlier reported results (Borch et al. 2026) of a syngas-enriched microbial community consisting of *Thermoanaerobacterium* and acetogens. *Thermoanaerobacterium* is a hydrolysing strain that produces H_2_:CO_2_ from amino acids/yeast extract (Dotzauer et al., n.d.; Sim et al. 2023; Jain et al. 2024). We hypothesize that the initial pressure increase is H_2_:CO_2_ from the hydrolysers, which is subsequently taken up by the acetogens. When syngas (H_2_:CO:CO_2_, 57:30:13, 2.5 bar) was the only substrate, a constant pressure was observed for the first 24 h, and then the pressure started decreasing exponentially, before eventually levelling off at a lower and constant pressure. When co-feeding fructose to syngas early, an initial gas production peak is observed, similar to when fructose is fed alone. However, when co-feeding fructose in the middle of the gas uptake phase, there are no changes in the gas pressure curve (Figure 4A). The fructose addition does at this point not change the acetogenic metabolism drastically away from gas-uptake, and it might even support ATP generation and additional biomass formation. The hydrolysers have a higher maximum growth rate than the acetogens, but the acetogens are either taking up the sugar, or if the hydrolysers begin growing and producing H_2_:CO_2_, this produced gas is then also taken up at an equal rate, as no change in the head-space pressure curve is seen. For the early fructose feeding (5 h after inoculation) the H_2_:CO_2_ production is also seen (Figure 4A). Because the pressure slope for this cultivation is less steep, the total gas uptake rate is slower. The fructose could have shifted the acetogen metabolism slightly away from gas, or the added fructose has potentially decreased the CO inhibitory effect, resulting in earlier onset of the H2/CO2 uptake, potentially in parallel with the CO. This aligns with other studies showing that co-feeding can limit the CO inhibition (Mann et al. 2020).

**Figure 4.**
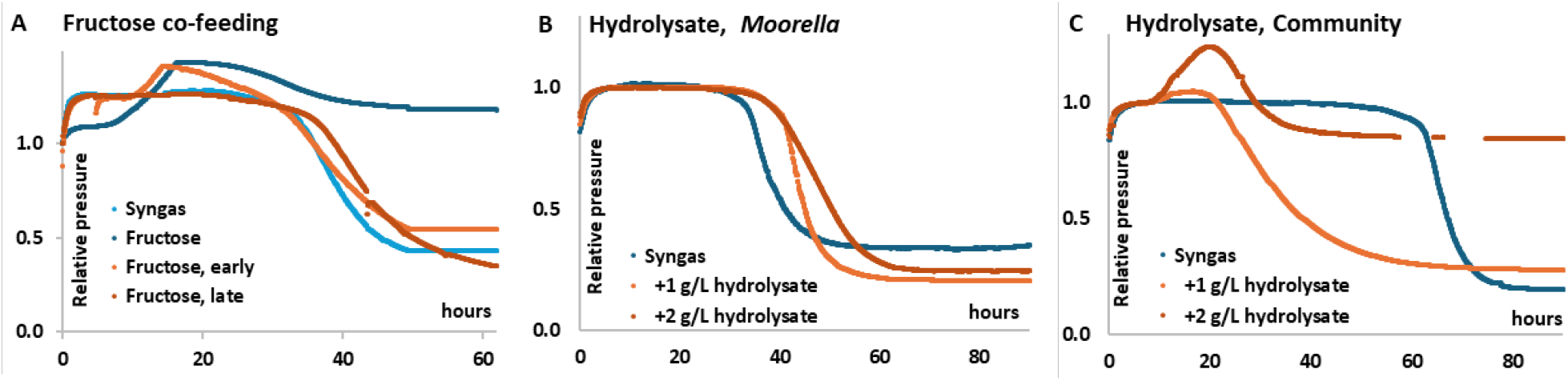
Pressure response when co-feed syngas fermentations. Pressure curves of co-feeding microbial communities and pure *Moorella* at various time points and substrate concentrations. A) Syngas fermenting microbial community co-feed fructose (1g/L) at various time points. B) *Moorella* syngas fermentation, co-feed hydrolysate at 1 or 2 g/L xylose. C) Syngas fermenting microbial community, co-feed hydrolysate at 1 or 2 g/L of xylose. The hydrolysate also contains small amounts of other sugars and lignocellulosic degradation products.

Next, wood-chip hydrolysate was co-fed with the syngas to the microbial community and a *Moorella* pure culture, at the beginning of the fermentations (Figure 4B,C). The hydrolysate mainly contained xylose with small amounts of other sugars and some soluble lignin degradation products. It was fed in amounts that resulted in a xylose concentration of either 1 or 2 g/L, thus resulting in a slightly higher total substrate concentration. For the microbial community, co-feeding hydrolysate shortened the initial phase with stationary pressure. There is a H_2_:CO_2_ production peak, as seen with fructose addition. However, this peak is not sharp; it transitions to re-uptake more gradually, indicating that gas uptake starts faster than for fructose and occurs before the hydrolysers have consumed all the xylose. When adding 2 g/L of hydrolysate, the gases are not completely re-consumed, this amount shifts the metabolism and the interspecies dynamics. For the pure *Moorella* culture, the hydrolysate addition did not result in a pressure increase peak as for the microbial community, supporting the hypothesis that this peak is created by the hydrolyser in the microbial community. Both 1 and 2 g/L hydrolysate additions to *Moorella* did not limit total gas uptake but, conversely, resulted in a lower final pressure, indicating that more electrons were used for CO_2_ reduction. Overall, the online pressure sensors could visualize the gas response kinetics of a syngas-fermenting community under various co-feeding strategies. When additional carbon (and electrons) are co-fed, the time and amount of the cofeeding seem crucial, and the variations result in different metabolic dynamics. To investigate this further, the resulting gas and product composition should be measured, and the carbon balances estimated.

### 4.3 Demonstrating a net zero CO_**2**_ **fermentation**

With the aim of continued investigation of microbial production of sustainable chemicals a new experiment was setup. Because gas fermentation is limited by H_2_ and CO mass transfer, we aimed to co-feed liquid substrates to achieve full conversion without substrate loss to CO_2_. We made a co-feeding experiment with formate and methanol. These substrates are liquid and fully soluble in water; they are single-carbon molecules (C1) and can be produced from CO_2_ waste stream sources. In the wood ljungdahl pathway (WLP), formate is directly part of the methyl branch or is either activated or oxidized to CO_2_ and integrated into the carbonyl branch. Methanol is linked to the WLP via a methyltransferase system that acts as a methyl-group donor, bypassing the reductive methyl branch. (Kremp & Müller, 2021). These substrates further eliminate the ATP-consuming step required for CO_2_ fixation, thereby improving overall energy retention (Kremp et al., 2018; Schuchmann & Müller, 2014). The right balance between highly reduced methanol and the less reduced formate can balance the supply of carbon and reducing equivalents, resulting in no net CO_2_ loss. To illustrate this, a mesophilic (30 °C) microbial community, enriched on methanol and formate from Danish soil was used.

We used online pressure sensors to monitor the cultivation. Based on the previous experiences, the substrates were added from the start, a 100% nitrogen headspace with 1 bar overpressure was applied, the vials were not sampled during the cultivation. The community was cultivated at five conditions. Two were formate and methanol individually, one with the combination, and two controls, a positive control with fructose, and a negative control with base medium and no substrate. The cultivations were performed in closed 120 ml vials containing 50 ml of medium, rubber stoppers, and a nitrogen headspace at 1 bar overpressure. They were done in minimum triplicate (n≥3). Through pressure measurement, biomass production (OD600), and product formation (HPLC) the carbon balances were estimated and titers and yields determined.

For the positive control, the glucose is consumed, sharply increasing the pressure to +42%, and then it remains constant (Figure 5A). It produces 33 cmmolL-1 acetate, 22 cmmolL-1 ethanol, and the highest biomass concentration 0.57 g_DW_L^-1^ (Figure 5B,C). The negative control, although minimal, showed some biomass growth (0.06 g_DW_L-1) and acetate production (9 cmmolL-1) due to the yeast extract in the medium, but no pressure change was observed. Methanol did not promote growth, and the pressure, biomass, and product formation were comparable to those of the negative control. With formate only, the pressure clearly increased (12%), but most of the substrate (83%) remained unconsumed, and the biomass (0.08 g_DW_L-1) and acetate formation (12 cmmolL-1) just barely exceeded the negative control. This indicated that formate is mainly oxidized to CO_2_ and does not provide ATP/energy for growth. In contrast, when co-feeding methanol and formate (methanol:formate) together, the pressure increases, comparable to the formate-only condition, but the pattern shifts (18h), and the pressure gradually decreases back to the initial pressure. Resulting in a cultivation with no pressure change, and thus no net loss of CO_2_, in contrast to the glucose and formate conditions. Additionally, 0.18 g_DW_L^-1^ of biomass is formed, significantly higher (p = 0.029; *t*-test, unequal variances) than for the formate-only condition. Acetate (32 cmmolL^-1^) was the only product, in a concentration similar to the glucose control. As no ethanol is formed, total product formation is lower than in the glucose control, but because biomass formation is lower, the product-to-biomass ratio is higher. The presence of methanol approximately doubles the formate consumption. The two pressure phases indicate that two separate processes are occurring. An initial formation of CO_2_ from formate, and a subsequent and slower reuptake of CO_2_, enabled by the electron and reducing power of methanol. It is thus not a coupled, balanced metabolic process but two uncoupled processes. It could be two different microbes working in parallel (Rotaru et al. 2012), or it could be one microbe, with formate dehydrogenase activity, and the ability to subsequently take up the CO_2_ and H_2_ formed (Schuchmann and Müller 2013). A substrate with a high electron-to-carbon ratio (e.g., methanol, glycerol) will require a co-substrate (electron acceptor) with an electron-to-carbon ratio lower than that of the biomass for growth to occur. Thus, no growth is observed on methanol before CO_2_ is present, which is generated from formate.

**Figure 5.**
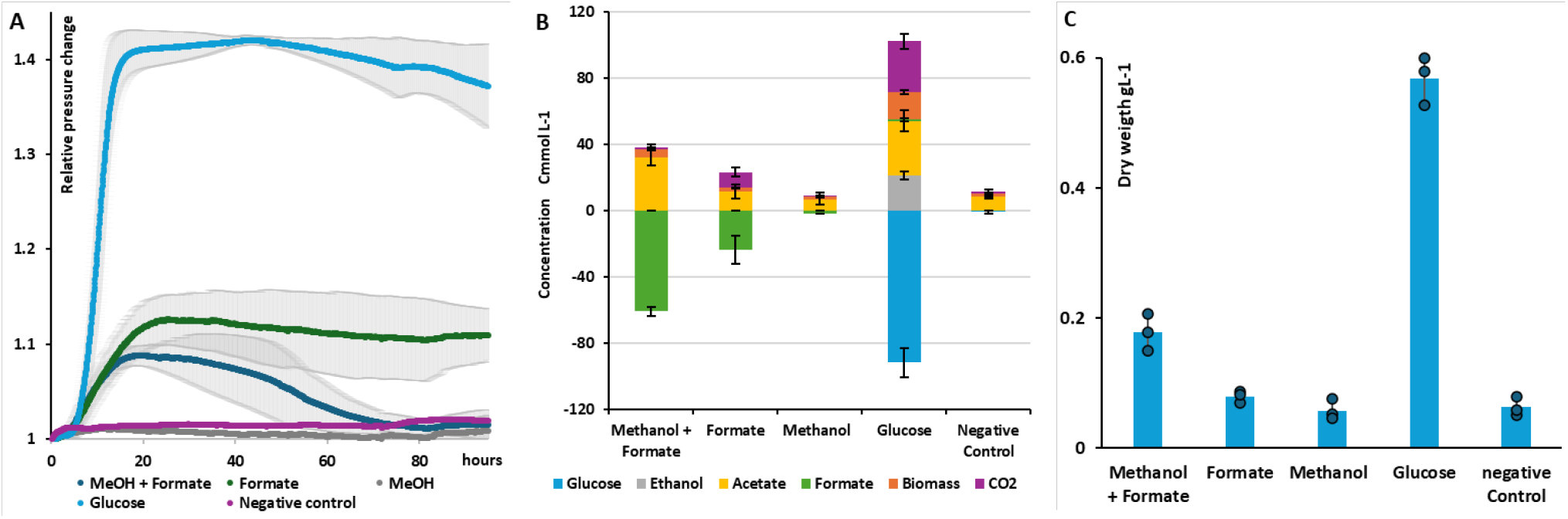
Co-feeding a methanol:formate-enriched microbial community. A) Resulting relative pressure curves of the co-feeding and control conditions. Starting headspace pressure 1 bar, 100% N_2_. B) Estimated carbon balance and product formation in cmmolL-1. Consumption is indicated by negative values. Methanol remained undetectable under the methods applied. C) Final dry weight (DW) gL-1. Converted from OD600 measurements at the end of cultivation (125h). All experiments were done in 120 ml anaerobic vials, with 50 mL medium, at 30 °C, and with magnetic stirring at 350 rpm. Substrates were added at the start of the cultivations at an equimolar concentration of 0.24 cmolL-1. The methanol:formate conditions were 0.12 cmolL-1 methanol and 0.12 cmolL-1 formate. Error bars indicate the standard deviation from independent triplicates or above (n≥3).

**Figure 6.**
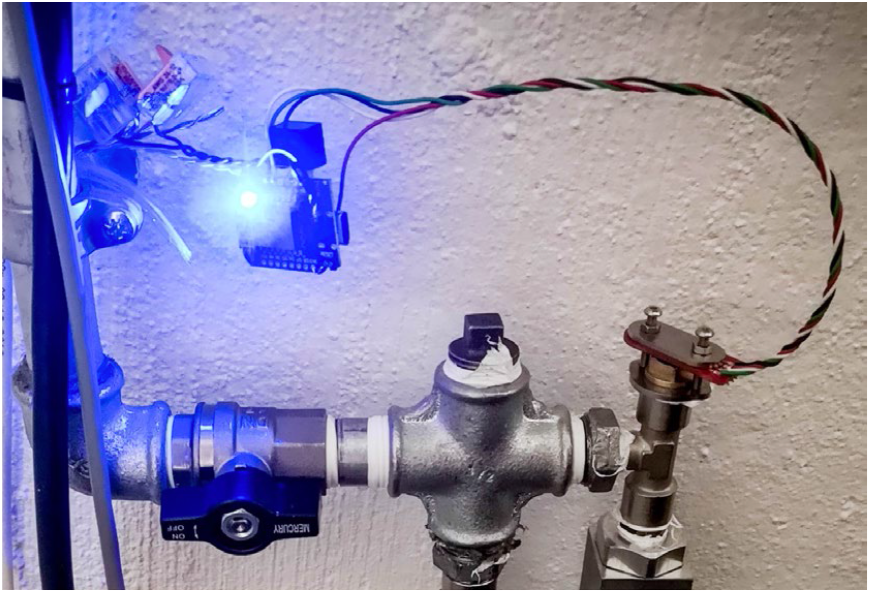
Compressed air monitoring. The sensor is used for extended monitoring of a 10 bar compressed air system. This setup employs a similar mounting system to the vial setup, except that the sensor is held against an O-ring on a pipe fitting, not a syringe. Additionally, a laser-cut stainless steel plate is added to stiffen the circuit board, as the measured pressure can reach up to 12 bar.

We successfully demonstrated a net-zero-CO_2_ fermentation with an enriched microbial community using a methanol:formate substrate combination. Both substrates were consumed, resulting in biomass and acetate production, with no final change in headspace pressure. This approach further demonstrates co-feeding as a method to circumvent gas mass-transfer limitations in gas fermentations. If an analogous process were implemented at an industrial scale, it could further reduce the need for tubing, pumping, and gas sparging. However, the pressure increases before it is taken up and should be considered in the reactor design.

To reach these results and conclusions, the online pressure sensors were used in small-scale batch cultivations. It was also assumed that the final pressure increases were due to CO_2_, and the residual gas for the methanol:formate condition was only the initial N_2_. This allowed for estimated carbon balances, and the gas kinetics further indicated a dual, non-coupled process. An anaerobic bioreactor sparged with N2 and CO2 and equipped with online gas analysis could have been used to reach the same conclusions, although it would require additional equipment and handling time. The culture is a microbial community, and multiple simultaneous gas reactions may occur, in particular, the production of methane and hydrogen. Thus, pressure alone does not provide enough information. For further application of the described tools and methodology, the final gas composition could be measured to validate the assumptions about the gas composition.

### 4.4 Online monitoring of compressed air

In addition to measuring gas pressure in vials, we have tested the sensor, breakout board and code (a modified version) for 24/7/365 monitoring of air pressure to various machines at Fablab RUC. In addition to improving lab reliability and safety, and protecting the machines from use under improper conditions, the setup has enabled us to evaluate sensor reliability. Two sensors have been running for 3 years without problems. This setup demonstrates the sensors’ robustness and the system’s viability for long-term, remote, online monitoring of experiments over extended periods. Testing also showed the importance of the extra checks and safety routines implemented in the code, including checking communication, validating data as sane, reinitialising sensors regularly, and rebooting the microcontroller if errors occur. If one does not take these steps, one occasionally gets readings that are way beyond believable ranges (e.g., 65000 or –65000 degrees or mBar), which makes it easy to discount them.

## 5 Discussion

### 5.1 Features and advantages

We designed and built an online pressure sensor system that met the initial design criteria. While a pressure sensor is not novel, we highlight the following key achievements, features, and advantages relative to prior art.

- It integrates with sterile and anaerobic small-scale laboratory batch cultivation workflows.
- The sensor resolution, quality control steps, and the potential of not disturbing experiments along the cultivation, provide unexpectedly high quality of online pressure data, exceeding some of the currently available laboratory systems.
- The sample rate and data quality allow for the detection of gas rates and, therefore, microbial catabolism in real-time.
- A high gas-tightness due to limited polymer fittings and tubing.
- The sensor is robust and operational even under humid conditions and at high temperatures.
- It is usable where high gas-tightness, over time, against pressure, is required
- It is compact and lightweight, allowing each sensor to be mounted on small laboratory flasks or vials without additional support, and its size also makes it possible to mount multiple sensors in the same space as vials or flasks that fit in laboratory incubators.
- Applying wires to power and reading the sensor provides long-term robustness (>3 years)
- A simple user interface and user experience.
- It’s affordable.
- An open-source science hardware solution that allows for future collaborations and integrations.
- The sensor is rigorously tested in quantities, across various use cases, and over an extended period.

### 5.2 Current limitations

The two main challenges identified through extensive user testing were that the sensor should not be moved, as doing so can cause 1) temperature fluctuations, which can lead to noisy data, and 2) leakage along the sides of the needle. Rotary shaking is problematic unless the sensor is properly fixed. Leakage is usually evident and can be considered during analysis. Worn-out rubber stoppers often cause leaks. Magnetic stirring provided reliable data and was used in the experiments described. Hydrogen does leak from the sensors, though in limited quantities. During long experiments, spanning several days, this should be considered in experimental design, with appropriate blank and control conditions, and kept in mind during data analysis. The sensors are factory calibrated, but we still see a +/- 10-50 mbar deviation between the sensors, though the individual relative sensor readings are accurate. Data should be averaged and normalised by measuring atmospheric pressure with all sensors before the experimental run. Pressure will gradually change until temperature and water vapour have equilibrated, as discussed in the biogas case section 4.1. It is advisable to ensure stable, similar readings before inoculating experiments, as this allows time to refit the sensor or repressurize the bottles, if needed. The sensors have proven incredibly robust, although some malfunctions and errors have been observed during extensive testing. Most commonly:

- The surface-mounted sensor breaks off the PCB during assembly or needle exchange.
- Connectors or wires break due to rough handling and bending.
- High humidity or spilling media directly on electronics can cause sensor read errors or complete malfunctions. These errors are filtered, and an error code is returned.
- Media enters the needle by inverting the vial or through horizontal or vigorous shaking. This can block the needle or disrupt the circuit, as mentioned above.
- Most errors are resolved by disconnecting the sensor, cleaning it, replacing the needle, inspecting the device, and drying the sensor.

### 5.3 Experimental design and data interpretation

In biological engineering, careful consideration is essential for designing experiments and analysing pressure data. Factors such as temperature differences between preparation and the actual experiment, as well as sparging with dry gas, can influence pressure measurements. It’s important to note that pressure will gradually change until temperature and water vapor reach equilibrium, as explained in the biogas results section 4.1. To ensure accuracy, allow the setup to equilibrate and confirm that pressure readings are stable before inoculating. This prevents mistaking microbial activity for temperature effects and provides quality control to verify consistent starting pressures. Whenever feasible, use dedicated incubators to minimize temperature fluctuations that could create noise in gas measurement data. Limit sampling to prevent gas leaks or temperature changes that could distort results. Incorporating blank and control conditions helps account for baseline gas leaks, particularly with high hydrogen levels. Avoid experiments involving shaking, as physical stress from shakers and shaking incubators can cause leaks, errors, or equipment failure. Instead, consider magnetic stirring or firmly mounting sensors and microcontrollers. Shaking can also cause medium to enter the needle, clog it, or short-circuit electronics. In screening experiments, an initial automated setup with more conditions can evaluate the conditions, timing, and manual sampling required for a subsequent run. The initial run can also help qualify what conditions make it into resource-limited, high-quality bioreactors. In microbial communities, pressure results reflect the combined activities of different processes. Pressure may stay stable when microbes produce and consume gases simultaneously or display growth curves of various strains with different metabolisms in successive phases. The catabolic and anabolic reactions relevant to the experiment should be considered. Since catabolism precedes anabolism and biomass growth, we often observe rapid pressure changes immediately after inoculation, especially with rich media.

### 5.4 Perspectives

We developed and validated an open-source online pressure sensor system designed for sterile anaerobic vial-scale cultivations. By combining a gas-tight needle interface on a custom breakout PCB with a microSD-logging, Wi-Fi-enabled ESP32 microcontroller, we enabled continuous pressure and temperature monitoring without manual sampling. The case studies show that high-resolution pressure measurements can be used to quantify gas production or uptake rates and to resolve kinetic phases and metabolic transitions that are difficult to capture with manual manometer measurements. The successful deployment of more than 130 units across multiple laboratories further demonstrates practical robustness and reproducibility. The application is broad, particularly where suspended solids prevent optical measurements or for solid-state fermentations. We hope to further improve the robustness and usability of the sensors. Complementary measurements of gas composition, or connecting online to pH, redox, or backscatter, could extend future use.

## Supporting information

Supplementary data for mmborch 2026 Pressure sensing for microbial batch cultivation

## 6 Nomenclature

Nomenclature

AMPTS: Automatic Methane Potential Test System
ATEX: Explosive atmospheres (ATEX directives/standards)
ATP: Adenosine triphosphate
BOD: Biochemical oxygen demand
BMP: Biochemical methane potential
CSV: Comma-separated values
ESP32: 32-bit microcontroller with Wi-Fi (Espressif Systems)
GC-TCD: Gas chromatography with thermal conductivity detector
H_2_: Hydrogen
HPLC: High-performance liquid chromatography
HTML: HyperText Markup Language
I/O: Input/output
I2C: Inter-integrated circuit (serial communication bus)
IDE: Integrated Development Environment
JSON: JavaScript Object Notation
MS5837-30BA: Absolute pressure sensor used in this study (TE Connectivity)
OSHWA: Open Source Hardware Association
PCB: Printed circuit board
RUC: Roskilde University
SMD: Surface-mounted device
SSID: Service set identifier (Wi-Fi network name)

Symbols and units

ΔGr: Gibbs free energy change of reaction
p(H_2_): Partial pressure of hydrogen
bar: Unit of pressure (10^5 Pa)
mbar: Millibar (10^-3 bar)
°C: Degrees Celsius
mL: Millilitre
mmol: Millimole
gDW L^-1^: Grams dry weight per litre
cmmol L^-1^: Carbon millimoles per litre
cmol L^-1^: Carbon moles per litre
rpm: Revolutions per minute

## 7 Acknowledgments

Thank you to all our great colleagues who have provided valuable insights and support in the development. Especially Daniel Vestergaard Davidsen for designing the PCB board and Nicolaj “DZL” Møbius for support with wiring and component design. The anaerobic experiments were performed by Parisa Ghofrani-Isfahani, Postdoc at DTU Chemical and Biochemical Engineering in the group of Professor Irini Angelidaki. Also, thank you to the people who have or are currently using our system for their valuable feedback and patience during the development process.

## 10 Author Contributions and information

Conceptualization, hardware and method development and first draft, MMB. Data acquisition and analysis, UJN, MMB, PK. Data interpretation, revision, comments, and approval, (all authors).

## 11 Supplementary Material

Supplementary Material are available at journal website and on github, https://github.com/nxdf/laerke.

## 12 Data Availability Statement

The Arduino code, design files for PCB and lasercut parts this study can be found in the “Laerke” repository on Github, https://github.com/nxdf/laerke.

