## Supplementary data for mmborch 2026 Pressure sensing for microbial batch cultivation for "Design and development of online pressure sensing for microbial batch cultivation"

### 1 Sensor validation

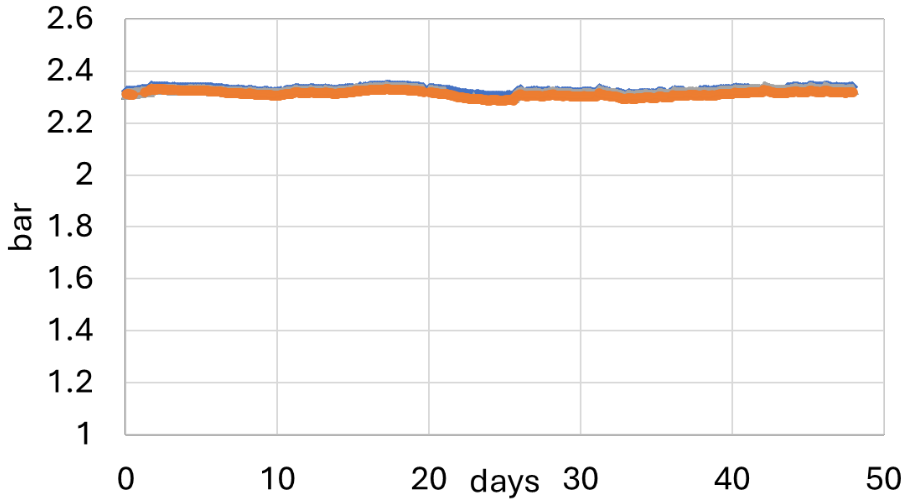


Supplementary Figure 1. Stable pressure reads for 1.5 months. Stable pressure in the vial with 1.5 bars overpressure of atmospheric air at fluctuating room temperature.

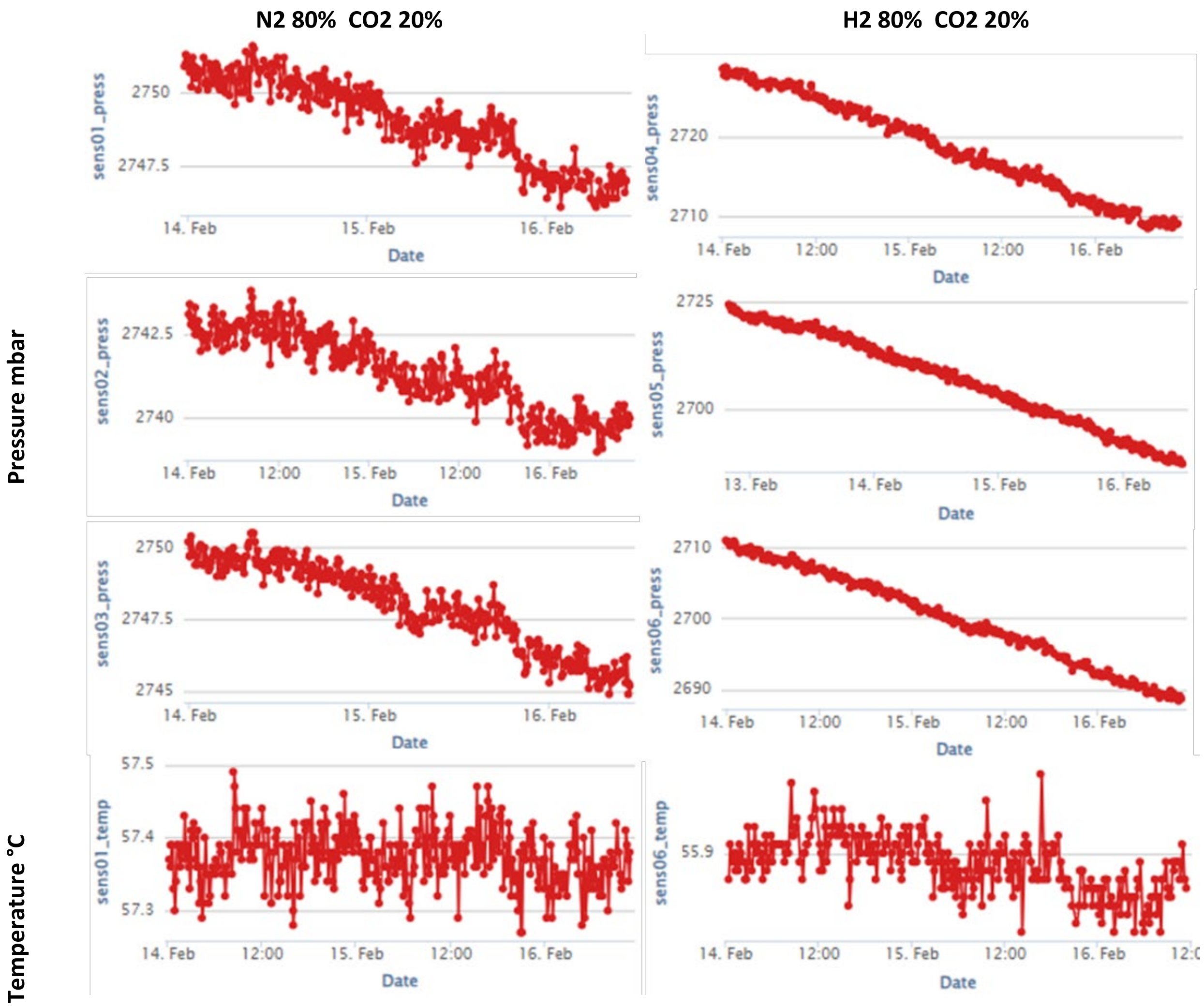


Supplementary Figure 2. Gas retention with overpressure of N2:CO2 80:20 or H2:CO2 80:20 in triplicates. Duplicate temperature reading.

### 2 Pressure sensor research

#### 2.1 The preexisting pressure sensor solutions

The existing pressure sensor solutions can be broadly categorized into three groups: 1) Commercial laboratory solutions, 2) Sensors designed for industrial-scale applications, and 3) Sensor breakout boards used for prototyping. They are described below; see supplementary materials for more details.

1) Gas composition experiments are typically done with manual sampling to analyze gas chromatograms for small batch cultures, but manual sampling has low throughput, and the equipment is expensive and not compatible with screening workflows. Pressurizable lab-scale stainless steel bioreactors are available, but they are usually operated at a fixed pressure with continuous gas flow. Lab-scale bioreactors are also very costly and require significant manual effort to operate. Systems for online pressure measurement in small-scale flask and vial batch cultures have been reported, especially for biogas monitoring (Fdz-Polanco et al., 2005; Himanshu et al., 2017; Pérez-Vidal et al., 2025). Additionally, commercial systems with pressure sensors mounted in the lid or cap are available, primarily for monitoring food spoilage, biogas production, yeast fermentations, biochemical oxygen demand (BOD), or bread dough rising (supplementary materials).

2) Pressure measurements are standard in fermentation and industrial chemical setups for process monitoring. For industrial use, they are usually high-quality, made of stainless steel, and designed to connect to process pipes and equipment, as well as integrate with data acquisition systems. These solutions typically cost around EUR 800 / USD 950 each. The sensors are often certified for use with explosive gases (ATEX). Modified with a luer lock fitting for sterile needle mounting, these sensors are frequently employed to sample batch cultures in laboratories. They are generally connected to a handheld display, from which pressure data is manually recorded in a logbook or data sheet.

3) There are many single sensors and several so-called “breakout boards" for different sensors, but none allow for gas-tight connections, or they are designed for tube connection, which could cause H2 leaks and potential O2 influx.

Although the issue has been encountered and resolved previously as reported in the literature, and some systems are available, none proved suitable for our use case. The high-quality industrial sensors were too expensive, heavy, and large to mount 20-45 of them inside a lab-scale incubator. The available pressure cap solutions were costly, did not fit the sterile, anaerobic vial cultivation setup with rubber stoppers, and had a pressure range that was too low for our needs. Additionally, the existing breakout boards lacked a form factor that allowed pressure-tight needle connections.

#### 2.2 Commercially available laboratory solutions

Bluesens yeast force:

- https://www.bluesens.com/us/products/gas-analyzers/bcp-series
- https://www.bluesens.com/fileadmin/user_upload/downloads-products/YeastForce/BlueSens%20Hefesensor%20YeastForce%2002-2021%20EN%20web.pdf

Ankom  Pressure sensor caps:

- https://www.ankom.com/product-catalog/ankom-rf-gas-production-system
- Lower resolution +-  2.8 mbar
- Lower accuracy:  +- 1 %
- \+ Pressure release possible
- Pressure range -10 + 500 PSI approx.  – 0.7 bar to + 34.4 bar
- 4750 USD for a complete setup with 5 caps.
- 32’500 DKK = 6500 pr bottle.
- Needs batteries
- hardware - Dedicated wifi module
- Dedicated software

OxiTop

- For biogas, BOD or respiration measurements.
- Pressure -500 – 1500 mbar
- https://www.xylem.com/en-us/campaigns/wtw-oxitop/

BPC intstruments, AMPTS®

- For biogas potential.
- https://bpcinstruments.com/bpc_products/ampts3/

Kuhner

- Oxygen transfer rates, online measurements.
- https://kuhner.com/en/products/data/Add-ons_KuhnerTOM.php

#### 2.3 Breakout board or single sensors

Sparkfun – DIY board

- Sensor with lower accurary
- Lots of examples available
- https://www.sparkfun.com/products/12909

MS5837 Breakout boards

- https://www.amazon.com/MS5837-Pressure-Sensor-Waterproof-Detectors/dp/B0CL59279G
- Same sensor mounted on a PCB.
- No resistors for connecting several sensors to the same controller, and no space for mounting anything gas-tight around the sensor.

NXP MPX4250AP - Farnell

- Analog piezoresistive transducer
- Absolute 20-250kPa – 2.5 bar
- Sensitivity 20mV / kPa
- 1.5% error
- Used in: Palucha 2025. bioresource technology 419. 132026

#### 2.4 Detailed code description

Full code on github.com/nxdf/laerke.

- Library from Bluerobotics that eases communication with the MS5837-30BA pressure sensor.
- Sensor Initialization & Reading: For each sensor, the code attempts initialization 5 times and reading 5 times. Upon successful read, it logs the sensor's pressure and temperature. If readings fail consistently, error messages are printed. The system also periodically resets power to sensors every two hours for reliability.
- System configuration parameters, including sensor pin assignments, network credentials, and reset intervals, are currently defined as static constants within the firmware. Future iterations of the system will implement a runtime configuration parser to load these settings directly from a config file on the SD card
- Data logging on the SD-card. Data logging occurs in the loop where sensor readings for pressure and temperature are retrieved sequentially from each sensor. These readings are appended to a CSV file on the SD card,
- The ESP device broadcasts a local WiFi Access Point using predefined SSID and password. It hosts a simple HTML page served via an embedded web server, allowing users to view live sensor data in a table, and to download the CSV log. The live data is pushed to the webpage using WebSockets using JSON format to update pressure and temperature readings in real-time.
- Function and sample rate. The standard setup reads the sensor every 3rd minute. The sensors allow for a reading approximately every 100 ms. Too high a communication rate to the sensor results in error values being returned, as the sensor needs to run an internal process and translate the data.
